# The cadmium-hypertolerant fern, *Athyrium yokoscense*, exhibits two cadmium stress mitigation strategies in the roots and the aerial parts

**DOI:** 10.1101/2023.12.06.570362

**Authors:** Yuko Ukai, Hiroki Taoka, Manaka Kamada, Yuko Wakui, Fumiyuki Goto, Kazuyoshi Kitazaki, Tomoko Abe, Akiko Hokura, Toshihiro Yoshihara, Hiroaki Shimada

## Abstract

*Athyrium yokoscense* is hypertolerant to cadmium (Cd) and can grow normally under a high Cd concentration despite Cd being a highly toxic heavy metal. To mitigate Cd stress in general plant species, Cd is promptly chelated with a thiol compound and is isolated into vacuoles. Generated active oxygen species (ROS) in the cytoplasm are removed by reduced glutathione. However, we found many differences in the countermeasures in *A. yokoscense*. Thiol compounds accumulated in the stele of the roots, although a long-term Cd exposure induced Cd accumulation in the aerial parts. Synchrotron radiation-based X-ray fluorescence (SR-XRF) analysis indicated that a large amount of Cd was localized in the cell walls of the roots. Overexpression of *AyNramp5a*, encoding a representative Fe and Mn transporter of *A. yokoscense*, increased both Cd uptake and iron and manganese uptake in rice calli under the Cd exposure conditions. Organic acids were abundantly detected in *A. yokoscense* roots. Investigating the chemical forms of the Cd molecules by X-ray absorption fine structure (XAFS) analysis detected many compounds with Cd-oxygen (Cd-O) binding in *A. yokoscense* roots, whereas in the aerial parts, the ratio of the compounds with Cd-sulfur (Cd-S) binding was increased. Together, our results imply that the strong Cd tolerance of *A. yokoscense* is an attribute of the following two mechanisms: Cd-O compound formation in the cell wall is a barrier to reduce Cd uptake into aerial parts. Thiol compounds in the region of root stele are involved in detoxication of Cd by formation of Cd-S compounds.

**Statements and Declarations:** No potential conflict of interest was reported by the authors.

## Introduction

Cadmium (Cd) is very toxic to living organisms including most plants (Clemens 2006). The toxic concentration of Cd in ordinary plants is generally 5–10 μg g^-1^ dry weight (DW) (White and Brown 2010). However, a few species can grow in soils containing much higher concentrations of Cd than the normal limitation and are called Cd-hypertolerant plants. One such species, *Athyrium yokoscense* is a widely distributed fern in eastern Asia (Van et al. 2006; Yoshihara et al. 2005). *A. yokoscense* shows normal growth features under culture conditions containing 1000 μM Cd and accumulation of 1.8–3.8 mg g^-1^ DW Cd in the plant body over one month (Yoshihara et al. 2005).

Even though Cd is a nonessential element for plant growth, Cd is considered to be erroneously absorbed into plant cells competitively via transporters, such as Nramp and IRT transporters, that are ordinarily involved in the absorption of essential metals such as iron (Fe) and manganese (Mn) (Clemens 2006). As a result, Cd inhibits the absorption of essential metals and induces a shortage of the counteracted essential metals. In addition, Cd sometimes displaces essential metals such as calcium in living cells and interacts with the functional groups of proteins. This occasionally leads to the modification of protein structures and disturbs the cellular redox status. Furthermore, Cd is commonly detoxified by thiol compounds (substances possessing an -SH group), glutathione and phytochelatin to form a complex to isolate Cd in vacuoles (Hasanuzzaman et al. 2017). However, this mechanism occasionally damages living cells against their native functions. In detail, glutathione reduces reactive oxygen species (ROS) generated by transition metals such as Fe and removes them in cells (Andresen and Küpper 2013; Pompella et al. 2003). The binding Cd to glutathione and/or phytochelatin results in an increase in ROS (Clemens 2006).

Many species of plants secrete specific substances from their roots, which change the features of metal ions and other elements around the root region and thereby their absorbability. For example, it is known that barley secretes mugineic acid to solubilize insoluble Fe (Takemoto et al. 1979). Wheat and soybean acquire resistance to aluminum by chelating it with citric acid, oxalic acid, and malic acid secreted from roots and lowering their absorbability (Yang et al. 2012). A similar mechanism has been suggested for tolerance to Cd absorption by secreted organic acids (Xie et al 2013). A cabbage variety showing Cd tolerance was reported to maintain higher concentrations of organic acids in leaves and roots than a Cd-sensitive variety (Sun et al. 2013).

To understand the mechanism of tolerance to heavy metals in plants, the chemical forms of the metal must be determined. Commonly, the chemical forms of heavy metals in extracts from cells (soluble components) are analyzed by HPLC combined with inductively coupled plasma-mass spectrometry (ICP–MS). To date, the binding of Cd to glutelin has been detected in rice grains (Suzuki et al 1997).

Recently, studies employing synchrotron radiation (SR) to biological samples have increased because the analysis using SR X-rays enables chemical speciation with minimal sample preparation (Sarret et al. 2002). High-energy X-ray-based SR may clarify the chemical form of heavy metals including cadmium in plant samples. X-ray absorption fine structure (XAFS) analysis enables the nondestructive analysis of Cd in plants and exhibits the Cd complexation (Fukuda et al. 2008). It has been reported that Indian mustard (*Brassica juncea*) accumulates substantial amounts of Cd, which is associated with a rapid accumulation of phytochelatins in the roots, where the majority of Cd is coordinated with sulfur ligands, probably as a Cd-S complex (Salt et al. 1995). *Arabidopsis halleri* spp. accumulates a large amount of Cd in trichomes, where the majority of Cd exists in the divalent state and is bound to the O and/or N ligands (Fukuda et al. 2008). As described above, the chemical forms of accumulated Cd are largely different depending on the plant species. These results suggest that the mechanism of Cd accumulation diverges depending on the plant species.

We have shown that *A. yokoscense* accumulated Cd in distal roots, but only a small amount of Cd was detected in proximal roots and shoots when exposed to 0.1 μM Cd for 24 hours. In the roots of *A. yokoscense*, no marked difference was observed in the amount of ROS upon exposure to 100 μM Cd. Transcriptome analysis detected little significant difference in the gene expression level in *A. yokoscense* cells exposed to 100 μM Cd for 24 hours (Ukai et al. 2020). In general, plants induce the expression of the stress-responsive gene clusters when placed under the Cd condition. (He et al. 2015; Xu et al. 2012). Our observations indicate that Cd stress minimally induces the major responses indicated above in *A. yokoscense*. Thus, it is strongly suggested that *A. yokoscense* has an unknown Cd stress tolerance mechanism that cannot be explained by any known mechanisms. In this study, we evaluated the contribution of thiol compounds and organic acids to Cd tolerance. We also elucidated chemical forms of Cd by X-ray absorption edge structure (XANES) analysis and determined the localization of Cd in the root tissues of *A. yokoscense* by X-ray microbeam utilizing SPring-8. Based on these results, we discussed the mechanism of Cd tolerance in *A. yokoscense*.

## Materials and Methods

### Plant materials and conditions of Cd exposure

Callus-derived homogeneous plantlets of the fern *A. yokoscense* were prepared under *in vitro* conditions (Ukai et al. 2020; Yoshihara et al. 2005). We used plants at the sporophyte stage that were redifferentiated from cultured cells. *A. yokoscense* plantlets were grown in a magenta box containing 1/2 × MS medium (1/2 diluted Murashige and Skoog medium supplemented with 30 g L^−1^ sucrose, 0.3 % gellan gum, and vitamins, 1 mg L^−1^ glycine, 50 mg L^-1^ myo-inositol, 0.25 mg L^−1^ nicotinic acid, 0.25 mg L^−1^ pyridoxin-HCl and 0.05 mg L^−1^ thiamine-HCl) (Murashige and Skoog, 1962). The culture room was set at 25 °C, and plants were illuminated in a 14 h/10 h (L/D) cycle. Tobacco (*Nicotiana tabacum* L. cv. Petit Havana SR1) plantlets, which was known to be sensitive to Cd stress, were used as a control and were grown under the same conditions as the *A. yokoscense* plantlets.

For the Cd exposure experiments, plantlets were precultured on 1/2 or1/4 diluted MS medium for one week and then transplanted onto the same concentration of MS medium supplemented with 0, 100, 200, and 2000 μM CdCl_2_. They were cultured under long-day conditions.

### Image analysis of accumulated thiol compounds

The accumulation of thiol compounds on the surfaces of cut sections of roots was observed as follows: *A. yokoscense* plants were incubated on a 1/4 diluted MS medium (1/4 MS medium) containing 0 or 100 μM CdCl_2_ for 24 hours. They were harvested and stained with monochlorobimane (mCBI, Thermo Fisher Scientific, Massachusetts, USA) according to the method of Kováčik et al. (2014). Roots of plants were sliced into 100 μm thick sections using a Tabletop Hand Microtome TH (Kenis, Osaka, Japan) and subjected to observation of the stained cells by an ECLIPSE 80i upright fluorescence microscope (NIKON, Tokyo, Japan). Regions stained with mCBI were detected by 490 nm fluorescence with an excitation wavelength of 394 nm. The fluorescence intensity of stained roots was quantified using ImageJ 1.48 (http://rsb.info.nih.gov/ij/).

### Measurement of thiol compounds in Cd-exposed plants

*A. yokoscense* seedlings were transplanted to 1/4 MS medium with/without 100 μM CdCl_2_ and incubated for 24 hours. The underground parts of the plants were collected and ground to paste in liquid nitrogen. Then, 1 mL of 0.2 N HCl per 100 mg of plant samples was added to them to extract the acid-soluble fractions. The extracted fraction was neutralized by the addition of 0.2 M NaH_2_PO_4_ (pH 5.6), whose amount corresponded to 1/10 times the amount of HCl used for acid extraction, and 0.2 M NaOH, which was 4/5 times the HCl amount. The resultant fraction was used for the measurement of the total glutathione concentration. In addition, 1 µL of 2-vinylpyridine (VPD) was added to 200 µL of the total glutathione concentration determination solution to precipitate the reduced glutathione. After centrifugation, the supernatant fraction was used for measurement of the oxidized glutathione.

Glutathione was measured by the recycling assay according to Queval and Noctor (2007). The method relies on the glutathione reductase-dependent reduction of 5’5-dithiobis (2-nitrobenzoic acid) (DTNB), monitored at 412 nm. To measure glutathiones aliquots of 10 μL extract were added to plate wells containing 180 µL reaction solution composed of 100 mM Na_2_PO_4_ (pH 7.5), 5 mM EDTA, 0.5 mM NADPH, 0.5 mM DTNB, and 0.2 unit of glutathione reductase. After 5 minutes of incubation, the change in absorbance at 412 nm was detected every 10 seconds 80 times using Multiskan GO Microplate Spectrophotometer (Thermo Fisher Scientific). As the controls, 100 μM GSH and GSSG (FUJIFILM Wako Pure Chemical Corp., Osaka, Japan) were prepared and used for calibration of these concentrations.

### Determination of Cd and Zn in *A. yokoscense* by ICP–OES

*A. yokoscense* plants were incubated on 1/2 MS medium containing 0, 200, or 2000 μM CdCl_2_ for 2 weeks. Plant samples were collected and washed with deionized water three times, dried, frozen in liquid nitrogen, and lyophilized. The resultant tissues were divided into aerial and underground parts. These samples were ground using a mortar and pestle, dried at 80 °C for 4 hours, and cooled down in a desiccator. One hundred milligrams of the sample were combined with 2 mL of concentrated HNO_3_ and mixed well overnight. After this treatment, the reaction cocktails were placed in a sealed container P-25 for microwave decomposition (San-Ai Kagaku Co Ltd., Nagoya, Japan) and subjected to microwave decomposition for 10 minutes. After evaporation on a clean bench, the resultant residues were dissolved in 1 M HNO_3_. After filtration through a PTFE membrane filter (0.45 μm), they were filled up to 50 mL by the addition of 1 M HNO_3_ and used as sample solutions. Yttrium (Y) was added to these solutions as an internal standard element at 1 ppm.

The concentrations of Cd in the digested solution were quantified by ICP– OES (SPS3520UV, Hitachi High-Tech Science Corp., Tokyo Japan). The amount of Zn was also determined. Quantification was performed by the calibration curve method (Carter et al., 2018) using calibration curves at 214.506 nm for Cd, 202.314 for Zn, and 371.030 nm for Y. For the analysis, the following conditions were adopted: the high frequency (RF) output was 1.2 kW, the carrier gas was Ar, the plasma gas flow rate was 16 L min^−1^, the auxiliary gas flow rate was 0.6 L min^−1^, and the photometric height was 12 mm. The accuracy of the measurement was evaluated using the reference materials, SRM 1570a spinach leaves and SRM 1573a tomato leaves (National Institute of Standard and Technology, NIST, USA).

### Preparation of the *AyNramp5a* gene, and detection of Cd in transformant rice calli

Based on the nucleotide sequence data of *A. yokoscense AyNramp5a* cDNA, which registered as Athyo15474 in DRA008924, a fragment corresponding to the coding region of *AyNramp5a* was chemically synthesized. This artificial *AyNramp5a* gene was modified to optimize the codon usage for rice cells, and a 6xHis tag was added to its 3’ end (Supplementary Fig. S2b). Using this fragment as the template the coding region for AyNramp5a-6×His was PCR-amplified using the primers attached to the attB1 and attB2 sequences on both sides. The amplified DNA fragment was introduced into pDONR207 (Thermo Fisher) by a BP clonase (Thermo Fisher) reaction. Then, the resultant fragment was transferred into pGWB2 (Nakagawa et al. 2009) by an LR clonase (Thermo Fisher) reaction to construct a binary plasmid containing the artificial AyNramp5a-6xHis gene (pGW2-AyNramp5a-6×His), which was expressed under the CaMV 35S promoter in rice transformant cells.

Transformation of rice, *Oryza sativa* L. cv. Nipponbare, was performed by the *Agrobacterium* method (Hiei et al., 1994). From the transformants, total RNA was prepared to confirm the expression of the transgene by semi-quantitative RT–PCR. The amplification was performed as follows: 94 °C for 5 minutes followed by 30 cycles, 94 C° for 30 sec, 55 C° for 30 sec and 72 C° for 30 sec. The primer sequences used were 5’-TTAATGGCGATGGTGCGACT-3’ and 5’-CCCGGTAAGCGGATTCTTGT-3’. The semi-quantitative value is the relative intensity of each band minus background, with PC as 1. ImageJ (http://imagej.nih.gov/ij, 1.53v) was used for this analysis (Supplementary Fig. S2c). Transformant calli showing semi-quantitative values greater than 0.2 were cultured in the N6D medium (Chu et al., 1975).

Transformant calli were cultured in R2 liquid medium (Ohira et al., 1973) for 24 hours, transplanted into the R2 liquid medium supplemented with 0, 10, and 100 µM CdCl_2,_ and incubated at 28 °C with shaking for 24 hours. After this treatment, the callus was washed with R2 liquid medium and lyophilized. The dried frozen cells were ground using a mixer mill (MM400, Retsch, Haan, Germany), and 30 mg of the crushed cells were used to make a tablet 10 mm in diameter using a manual hydraulic press (GS15011; Specac Ltd., Orpington, UK) at 5 tons of pressure for 3 minutes. Transformant cells independently generated were determined the expression of the transgene. For usage of the triplicate analysis, transformant lines whose expression of the transgene had been confirmed were cultured. Then they were combined and used to prepare a tablet. This operation was performed three times and obtained three tablets in total. The obtained tablets were provided for the analysis of Cd and other elements by an energy dispersive X-ray fluorescence (XRF) spectrometer (Epsilon 5; Malvern Panalytical, Spectris, The Netherlands).

### Analysis of organic acid composition in roots

*A. yokoscense* was cultured for 0 and 14 days on 1/4 MS medium containing 0.4 % gellan gum (FUJIFILM Wako Pure Chemical Corp., Osaka, Japan) supplemented with 0 or 100 µM CdCl_2_. After harvesting, their root sections were excised and used for the preparation of the fractions of organic acids according to the method of Jonson et al. (1996): The root sections were ground in liquid nitrogen, weighed, added to 2 % HCl, and heated at 80 °C for 10 minutes. After the removal of precipitates by centrifugation, the resultant supernatant was extracted with chloroform to obtain the hydrophilic fraction that contained compounds of organic acids. Similarly, tobacco plants were exposed to Cd, and root fractions were prepared. They were provided to analyze the representative organic acids.

The amounts of organic acids were analyzed using an ACQUITY UPLC system (Waters Corp., Milford, MA, USA.). Organic acid compounds were separated using a separation column, ACQUITY UPLC HSS T3, with particle size: 1.8 µm, inner diameter: 2.1 mm x length 150 mm (Waters Corp.) at a column temperature of 30 °C and a pressure of 12000 psi. The organic acid compounds were eluted in 5 mM NaH_2_PO_4_ (pH 2.8) and detected using a PDA eλ detector (Waters Corp.) at 210 nm.

As the standard solutions, 10 mg mL^−1^ solutions of citric acid, sodium acetate, maleic acid, DL-isocitric acid, succinic acid, shikimic acid, oxalic acid, L-lactic acid, phytic acid, fumaric acid, and D-malic acid and 125 mg mL^−1^ pyruvic acid solution were used. Using the standard solutions, the calibration curve was produced. Fractions prepared from plant tissues were diluted 10-fold with 5 mM NaH_2_PO_4_ (pH 2.8) and subjected to UPLC analysis. These samples were filtered with a 0.20 µm membrane filter right before analysis.

### Synchrotron radiation-based X-ray fluorescence (SR-XRF) analysis

X-ray fluorescence (XRF) is an elemental analysis technique that is mainly used to provide elemental infomation with material analysis such as metals and ceramics. In this study, the XRF analysis technique was applied to biological samples. SR-XRF is a technique that can analyze extremely small areas with micrometer resolution (Nakai 1997). The SR-XRF analysis was performed as described previously (Kashiwabara et al. 2021; Harada et al. 2010). The roots and stems of *A. yokoscense* seedlings were cut from the plants, embedded in an OCT compound (Sakura Tissue-Tek, Tokyo, Japan), and frozen using dry ice. Transverse sections were cut to a thickness of 30 μm (roots) using a 35A stainless-steel blade (Feather, Osaka, Japan) attached to a cryomicrotome (Leica 3050S, Wetzlar, Germany) at –20 °C. The samples were placed on a thin polyester film (Mylar film, Chemplex, Palm City, FL, USA) stretched over a 4 × 4 cm^2^ acrylic plate with a 0.3 diameter hole (Hokura and Harada 2017).

Micro-XRF measurements were performed on beam line 37XU at SPring-8 (Hyogo, Japan, Terada et al. 2004). The analysis conditions were described in Fukuda et al. (2008). The X-ray beam with an energy of approximately 30 keV was focused to 0.65 μm (horizontal) and 1.1 μm (vertical) using a Kirkpatrick-Baez mirror system, and the samples were cooled with liquid nitrogen sprayed by a low-temperature system (Rigaku GN2, Tokyo, Japan) during measurements to minimize any potential beam damage.

XRF is detected the emission of characteristic secondary X-rays from a material that has been excited by being bombarded with high-energy X-rays. Fluorescent X-rays from the sample were detected by a Si(Li)-solid-state detector (SSD) or silicon drift detector (SDD). An energy range (region of interest, ROI) was set for the X-ray fluorescent peak of each element, and the peak area obtained was used as the X-ray fluorescence intensity. Therefore, if the sample does not contain the target element, an image like a white noise signal is obtained. Images of the elements in the tissues were obtained by scanning both vertically and horizontally. The image data were normalized with respect to the fluorescent X-ray intensity and expressed on a 256-step color scale with the strongest point in red and the lowest point in blue to obtain a two-dimensional elemental distribution map.

### Chemical forms of Cd

The XANES region is sensitive to the electronic structure information of atoms and can be used to qualitatively assign the coordination geometry of central atoms, formal oxidation states, etc. XANES provides information about the valence of the absorption elements and crystal structure of specimens (Kato et al. 2008). The comparison of the cadmium L_3_-edge XANES spectra of the model compounds showed that this technique enables a clear distinction between S and O ligands (Isaure et al. 2006). The XANES analysis of the Cd K-edge absorption edge spectra of Cd-S and Cd-O was performed as described previously (Fukuda et al. 2008; Yamaoka et al. 2010). Four samples were prepared from each of the aerial and underground parts of Cd-treated plants. They were frozen with liquid nitrogen, lyophilized, and powdered by pestle and mortar. Tablets were formed by pressing 100 mg of tissue powder at a pressure of 6 ton f cm^-2^ for 5 minutes using a 10 mm diameter tablet forming machine. This tablet was sealed with Mylar film and set on an acrylic plate with a hole of 2 cm diameter. The cadmium K-edge XAFS analyses were performed at the NW10A of PF-AR in the High Energy Accelerator Research Organization, Japan. The reference chemicals were CdO (Kanto Chemicals, Tokyo, Japan), CdS, Cd(NO_3_)_2·_4H_2_O, CdCl_2_, Cd(CH_3_COO)_2_, CdSO_4_, Cd-pectin, Cd-phytochelatin and Cd-methionine (Sigma 9038-94-2). Cd-pectin and Cd-phytochelatin were prepared according to Isaure et al. (2006) and Salt et al. (1995). XAFS spectra were analyzed using REX 2000 program ver. 2.57 (Rigaku, Tokyo, Japan). The obtained spectra were smoothed by the Savitzky-Golya method (Savitzky and Golya 1964). The mixing ratio of Cd-O and Cd-S was obtained according to the method of Isaure et al. (2015). Means of the values of the mixing ratios in each part were taken as and the quantitative ratios of Cd-S and Cd-O. The reliability factor for the goodness of fit, *R* factor, was calculated as residual between fit and experimental data.

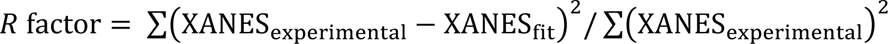

## Results

### Thiol compounds were generated in root stele due to Cd stress

Glutathione compounds consist of reduced and oxidized glutathione (GSH and GSSG). The ratio of the amount of GSSG is known to increase in response to environmental stress. We measured the amount of glutathione compounds in *A*. *yokoscense* to determine the physiological response to Cd stress. A decrease in the total amount of glutathione compounds was observed when the plant was exposed to 100 μM Cd (Fig. 1a). However, the concentration of GSSG was estimated to be very low due to being below the lower limit of detection (Fig. 1a). This fact indicates that most of the glutathione compounds were GSH.

**Fig. 1.**
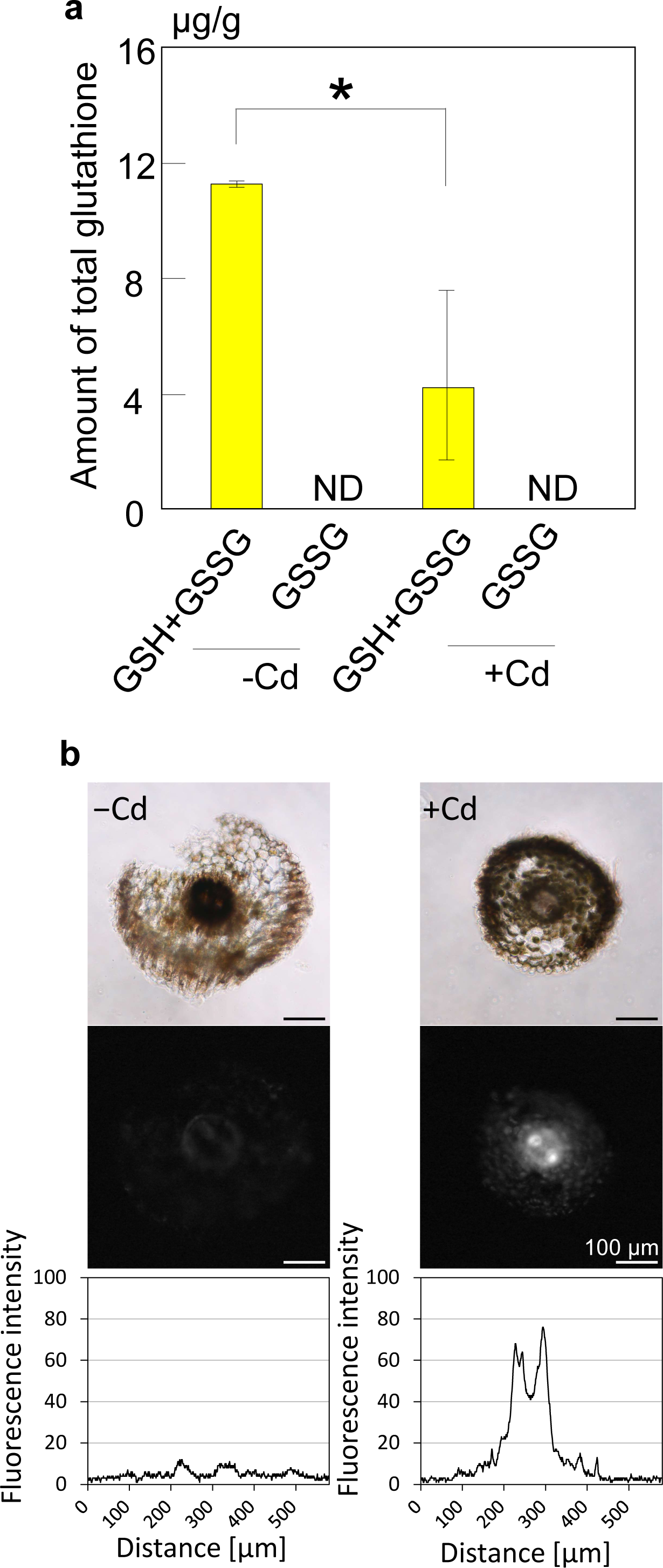
(a) Changes in the amounts of total and oxidized glutathione. GSH+GSSG: amount of total glutathione containing the oxidized GSH and reduced GSH, GSSG: amount of the oxidized glutathione. +Cd and -Cd indicate the results of the amount in the cells with/without treatment of 100 μM CdCl_2_ (*n* = 4). The asterisk indicates statistical significance by *t* test (*p* < 0.05). ND: not detected due to those below the limit of detection. (b) Detection of thiol compounds in roots. Cross-sections of the *A. yokoscense* roots were stained with mCBI. Upper and middle panels show the bright field and fluorescence images, respectively, with/without exposure to 100 µM CdCl_2_ for 24 hours. Lower panels show graphical images of the fluorescence intensity on an arbitrary line through the center of each root. Detection was performed at 394 nm/490 nm (wavelength of excitation/emission). Scale bars = 100 µm

Next, we analyzed the localization of accumulated thiol compounds in *A*. *yokoscense* treated with Cd stress for 24 hours. Some thiol compounds, such as phytochelatin and reduced glutathione, are known to function in an ROS scavenging system. Thiol compounds were detected in the stele of the root that was exposed to 100 μM Cd (Fig. 1b). Since no thiol compounds were detected in the roots of the plantlet without Cd exposure, they primarily accumulated in the stele of roots due to Cd exposure.

### Determination of Cd taken up into plants

To determine the behavior of Cd uptake in plantlets with long Cd-exposure, sporophyte-stage plantlets of *A. yokoscense* that were redifferentiated from the cultured cells were grown in 1/2 MS medium supplemented with 0, 200, and 2000 μM Cd for 2 weeks. These plants showed normal growth under these culture conditions. Cd accumulation in their aerial and underground parts was analyzed by an inductively coupled plasma optical emission spectrometer (ICP–OES). When they were cultured under 200 μM Cd conditions, Cd accumulated 337 and 207 μg g^−1^ (DW) in the underground and aerial parts of *A. yokoscense* sporophytes, respectively. Under the condition of 2000 μM Cd concentration, they reached 4880 and 3430 μg g^−1^ (DW) in the underground and aerial parts, respectively. Significant differences in Cd accumulation were found between underground and aerial parts (*p* < 0.05), whereas no significant alteration was detected in the amounts of Zn accumulation (Table 1). These results indicated that a large amount of Cd was absorbed and accumulated in the aerial parts depending on the Cd concentration in the medium when *A. yokoscense* was exposed to Cd for a long period of time. However, the amounts of accumulated Cd in the aerial parts were approximately two-thirds of those in the underground parts, suggesting that Cd transfer into the aerial parts was restricted.

**Table 1.** Cd accumulation in underground and aerial parts of the Cd-stressed *Athyrium yokoscense* (µg g^-1^ DW)

| Cd in the medium | Cd |  | *p*<br>value | Zn |  | *p*<br>value |
| --- | --- | --- | --- | --- | --- | --- |
|  | Underground | Aerial |  | Underground | Aerial |  |
| 0 $\mu\text{M}$ | ND | ND | — | $28.7 \pm 6.4$ | $31.9 \pm 4.9$ | 0.60 |
| 200 $\mu\text{M}$ | $337 \pm 30$ | $207 \pm 39$ | 0.020 | $26.5 \pm 0.9$ | $26.2 \pm 2.1$ | 0.88 |
| 2000 $\mu\text{M}$ | $4880 \pm 48$ | $3430 \pm 120$ | 0.015 | $29.9 \pm 1.0$ | $27.1 \pm 0.2$ | 0.054 |

Averaged values are indicated with standard deviation ( $n = 3$ ). ND indicates the value below the detection limit. The *p* values were calculated for the null hypothesis that there is no significant difference in the amount of Cd or Zn between the underground and aerial parts.

### *A*. *yokoscense* Nramp5 takes up Cd into rice transformant cells

OsNramp5 is Nramp family transporter located on the cell membrane of *Oryza sativa*, which is involved in Mn and Fe transport in their roots. We have reported the cDNA dataset that was established by RNA-seq analysis of *A*. *yokoscense* cells (Ukai et al. 2020). This report has revealed that *A*. *yokoscense* has eight genes encoding proteins showing high similarity to rice OsNramp5 (accession number Q8H4H5.1). These genes were named *AyNramp5a* to *AyNramp5h* according to the ratio of similarity to *OsNramp5*. Among them, the *AyNramp5a* (registered as *Athyo15474* in the dataset) and *AyNramp5f* (*Athyo44278*) genes were abundantly expressed in roots (Supplementary Fig. S1). We selected *AyNramp5a* showing 66 % similarity to *OsNramp5* (66.0 %) as the representative of *A*. *yokoscense* Nramp5 genes and used it for further analysis.

We constructed an artificial gene for AyNramp5a driven by the CaMV-35S promoter for overexpression in rice cali (Fig. 2a). Uptake of Cd, Fe, and Mn was measured in the resultant transformant callus that was cultured for 24 hours in medium containing 0, 10 and 100 µM Cd. The transformant callus showed increased uptake of Fe and Mn compared with the nontransformed callus, showing that AyNramp5a functioned as a Fe–Mn transporter. These amounts did not change significantly with the addition of Cd to the medium (Fig. 2b). The amount of Cd in the transformant was significantly increased due to the Cd concentration in the medium (Fig. 2b). These results indicate that AyNramp5a is involved in Cd uptake and still works well for Fe and Mn uptake under the Cd stress.

**Fig. 2.**
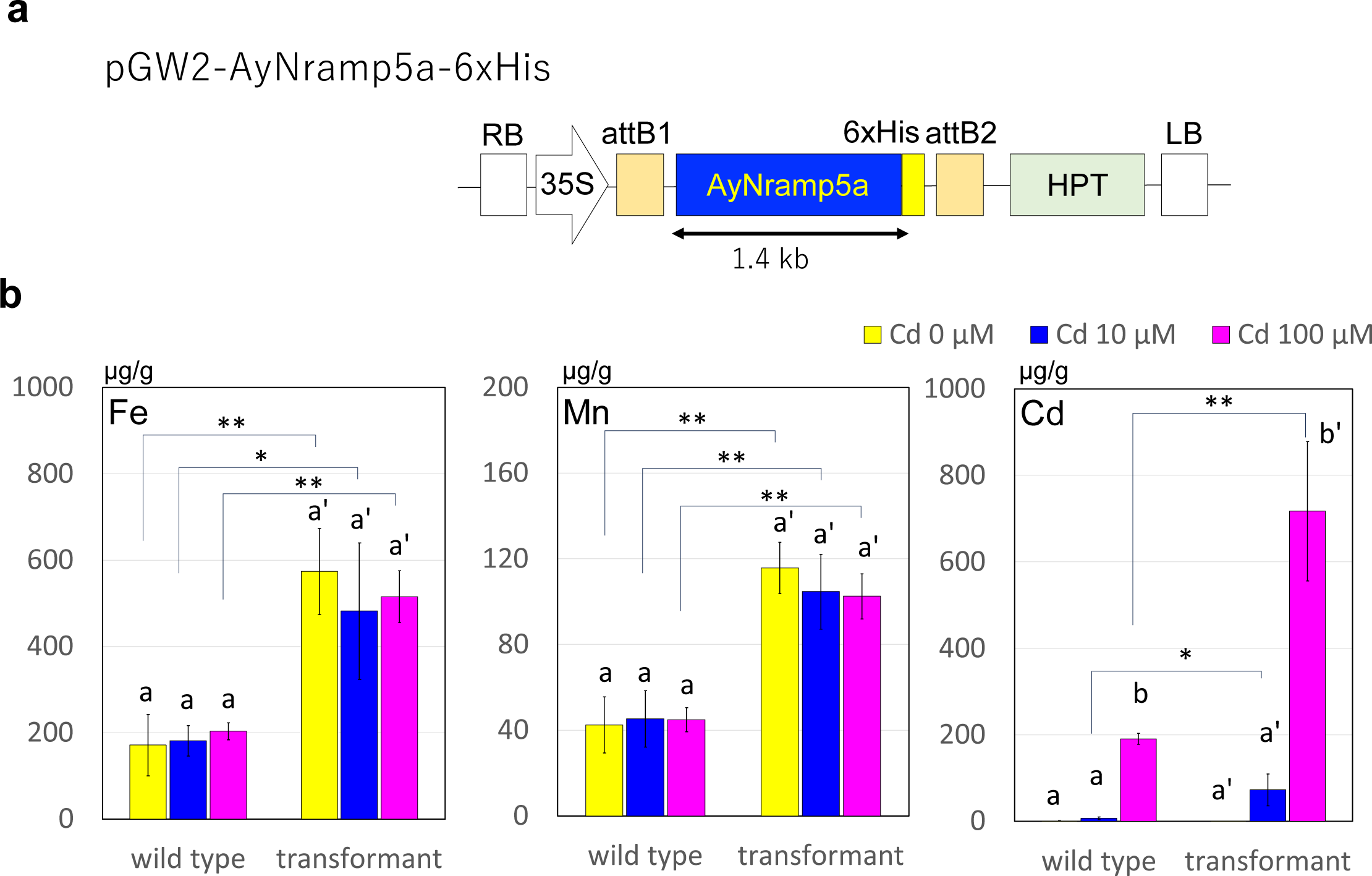
Detection of uptake of Fe, Mn, and Cd by *A. yokoscense* Nramp5a in the rice transformant callus. (a) Structure of pGW2-AyNramp5a-6×His, which is the artificial gene expressed in the rice callus. The nucleotide sequence of *AyNramp5a* is shown in Supplementary Figure S2. 6xHis: 6-fold His tag region. RB and, LB: Right border and Left border. 35S: CaMV 35S promoter. attB1 and attB2: sequences of attB1 and attB2, HPT: hygromycin resistance gene. (b) Uptake of Fe, Mn, and Cd in the wild-type and the transformant rice callus exposed to 0, 10 and 100 μM CdCl_2_ in the medium. In each graph, the amounts of the element in the rice callus were shown. The yellow, dark blue and magenta bars show, respectively the amounts in the callus exposed to 0, 10, and 100 µM CdCl_2_. Asterisks indicate the significant difference between wild type and transformant by *t* test (*n* = 3, \**p* < 0.05, \*\**p* < 0.01). Significant alterations in the amount of uptake depending on the Cd concentration were determined by the Tukey’s test at the 5% significance level.

### *A. yokoscense* roots accumulate large amounts of organic acids

Organic acids secreted from roots are known to immobilize/mobilize the metal ions. We analyzed the amounts of the representative organic acids L-lactate, D-fumarate, and D-malate in the *A. yokoscense* roots. A quantitative comparison of these organic acids revealed that the amounts of these organic acids were significantly greater in *A. yokoscense* than in tobacco (Fig. 3). This matter suggests that *A. yokoscense* root cells constitutively produced a large amount of these organic acid compounds.

**Fig. 3.**
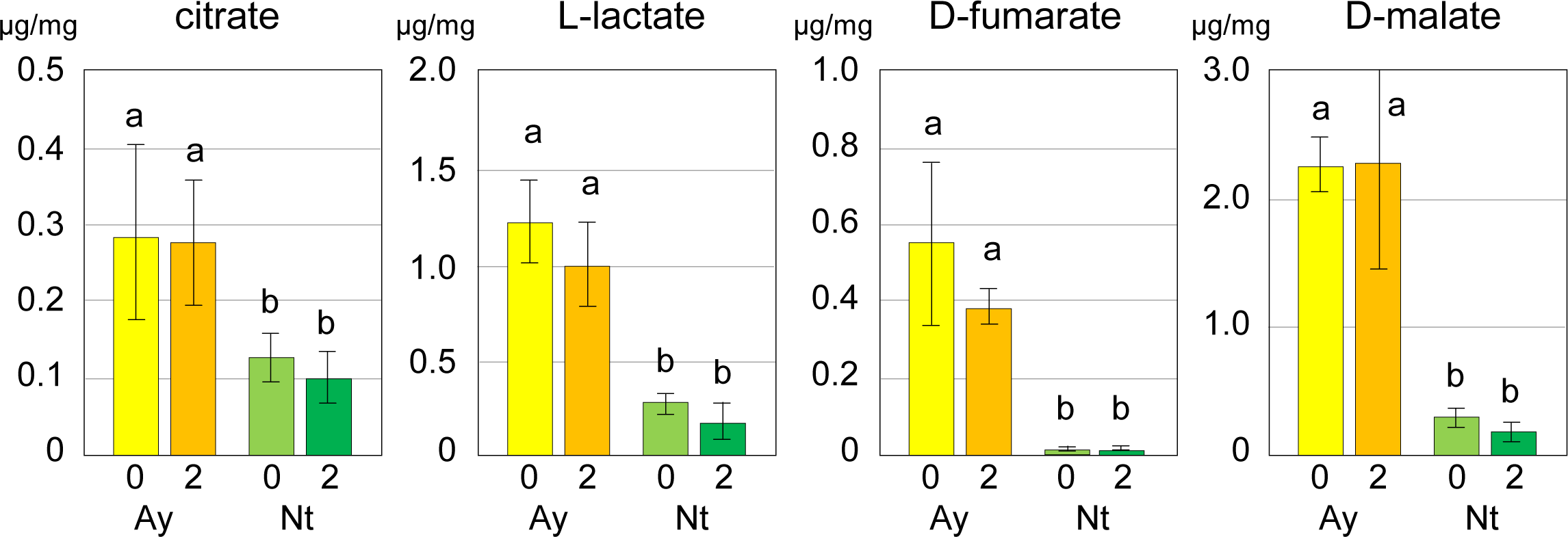
Amounts of representative organic acids in *A. yokoscense* and tobacco. The contents of intracellular citrate, L-lactate, D-fumarate, and D-malate are shown. Panels show the comparison of the contents of *A. yokoscense* (Ay) and tobacco (Nt). “0” and “2” indicate the values determined at 0 and 2 weeks after Cd treatment. Significant differences were detected by Tukey’s test at the 5% significance level. Yellow and yellow–green rectangles indicate the values of the non-Cd-treated *A. yokoscense* and tobacco cells, and orange and dark green rectangles indicate those of 100 μM CdCl_2_-treated *A. yokoscense* and tobacco cells. Bars indicate the standard deviations.

The amount of these organic acids showed no significant difference between 0 and 2 weeks after the beginning of Cd exposure in the roots of both *A. yokoscense* and tobacco (Fig. 3). This suggests that no induction of organic acid production occurred due to Cd exposure. This also shows that a large amount of these organic acids was still maintained in *A. yokoscense* root cells under Cd stress conditions.

### Cd is distributed primarily in the cell wall of the root tissue of *A*. *yokoscense*

In some plants showing relatively high resistance to Cd stress, it has been determined that Cd is abundantly distributed in the cell wall components of roots (Loix et al. 2017; Sterckeman and Thomine 2020). X-ray fluorescence (XRF) analysis using synchrotron radiation (SR) X-ray microbeam on frozen sections of 30 μm thickness visualized the distribution of Cd in the roots. In parallel, Zn, Fe, Mn, and K were analyzed as controls.

In the plants grown under 200 μM Cd conditions for 2 weeks, Cd was detected in the epidermis, cortex, endodermis, and stele (Fig. 4a). Zn, Fe, Mn, and K distributed in all tissues (Fig. 4a). Comparison with the microscopic images indicated that Cd was distributed mostly in the region corresponding to the cell wall and the intercellular space, and also accumulated in root vascular bundles (Fig. 4a); similar distribution was obtained in the plants grown under 2000 μM Cd exposure conditions (Fig. 4b). These results indicated that absorbed Cd was mainly localized in the root cell wall in *A*. *yokoscense*.

**Fig. 4.**
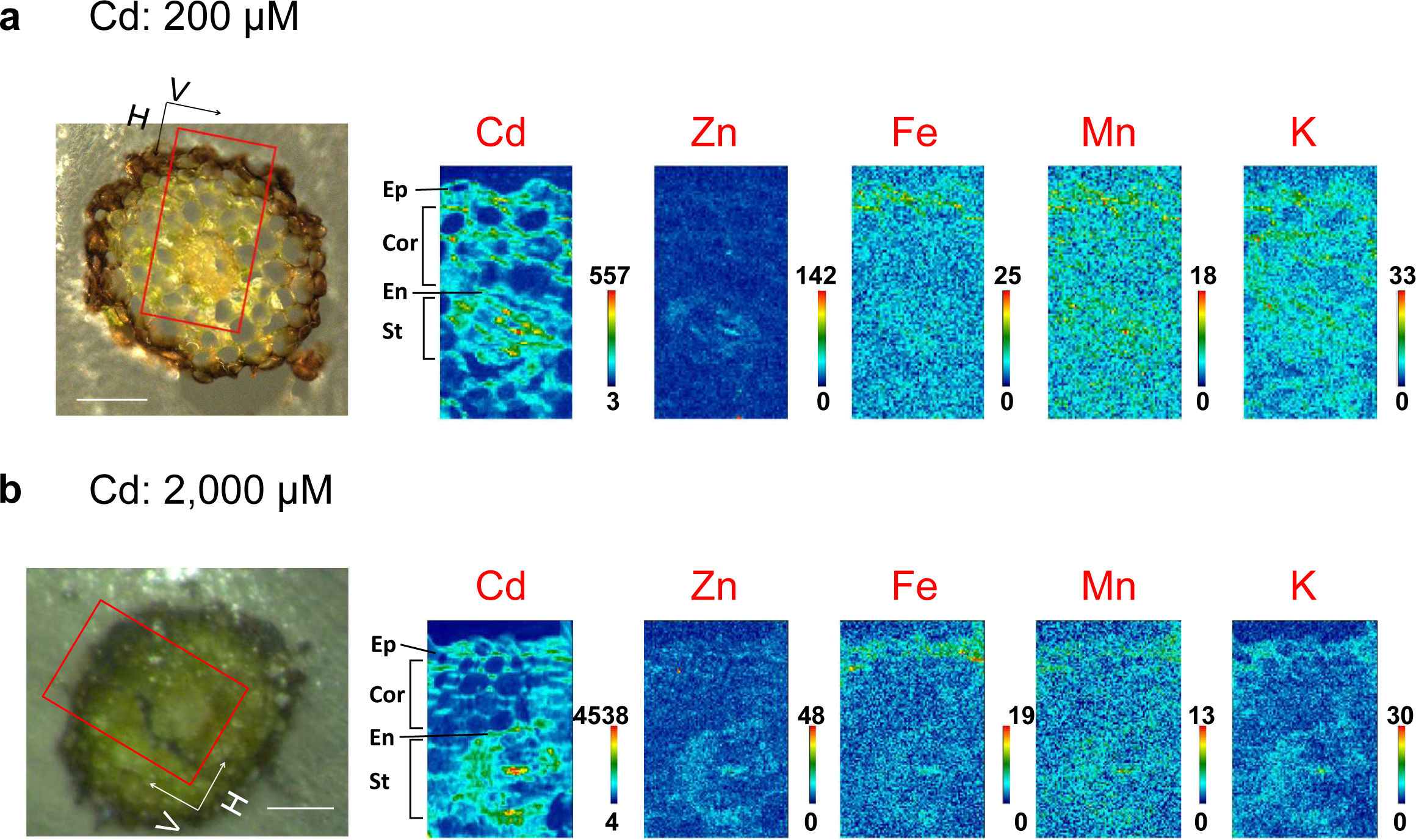
Distribution of elements in cross sections of root segments exposed to (a) 200 and (b) 2000 μM CdCl_2_. The left panels show the micrographs of *A. yokoscense* sporophyte roots. Right panels indicate the images of XRF analysis. Red squares indicate the measurement area shown on the right panels. They show the distribution of Cd, Zn, Fe, Mn and K, respectively. Color bars on the right indicate the relative values of the intensity of the signals of the corresponding elements. Detailed analytical conditions are described in Supplementary Table 1. Ep: Epidermis, Cor: Cortex, En: Endodermis, St: Stele, Scale bars = 100 µm

### Chemical forms of Cd accumulated in the underground and the aerial parts of *A*. *yokoscense*

The chemical form of Cd in the roots and the shoots was determined by X-ray absorption near-edge structure (XANES) analysis. The chemical form of Cd in the underground and the aerial parts can be estimated by comparing it with reference substances whose chemical form of Cd has been determined. For comparative verification, we used the reference substances involving a bond between Cd and oxygen, designated Cd-O, and those that included a bond between Cd and sulfur, designated Cd-S. To determine Cd forms in the fern, the samples were prepared from roots and aerial parts of sporophyte stage *A. yokoscense* that had been cultured under 200 µM and 2000 µM Cd. The electronic state of Cd is affected by the neighboring atom coordinated to Cd and exhibits a specific spectral shape showing patterns depending on the chemical form, leading to a clear distinction of whether it is attributed to the compound bound to oxygen or sulfur. When the sample contained both compounds of the Cd-O type and Cd-S type, the spectrum indicated the shape of the mixed feature according to the abundance ratio. Therefore, the quantitative ratio of Cd-O and Cd-S can be determined by pattern-fitting the XANES spectra using spectral data of the reference compounds (Salt et al. 1995; Yamaoka et al. 2010).

In the plantlet grown for 2 weeks under 200 µM Cd, the ratios of Cd-O and Cd-S were estimated to be 91 % and 9 %, respectively, in roots (Table 2). This indicates that Cd-O is the predominant chemical form of Cd accumulated in roots. In contrast, the proportions of Cd-O and Cd-S in the aerial parts were 59 % and 41 %, respectively, showing that the compounds with Cd-S bonds were more abundant than those in roots (Table 2).

**Table 2.** Ratio of chemical forms of Cd-oxide (Cd-O) and Cd-sulfide (Cd-S) accumulated in sporophyte stage plantlets of *Athyrium yokoscense* (%)

| Cd in the medium | parts | Ratio of Cd form |  | *R* factor <sup>a)</sup> |
| --- | --- | --- | --- | --- |
|  |  | Cd-O | Cd-S |  |
| 200 $\mu\text{M}$ | underground | 91 | 9 | 0.007 |
|  | aerial | 59 | 41 | 0.013 |
| 2000 $\mu\text{M}$ | underground | 72 | 28 | 0.012 |
|  | aerial | 64 | 36 | 0.010 |

Chemical forms of Cd in the plantlet were determined by XANES analysis. The sporophyte of *Athyrium yokoscense* was exposed to 200 $\mu\text{M}$ and 2,000 $\mu\text{M}$ $\text{CdCl}_2$ for 2 weeks. The ratios of Cd-O and Cd-S (%) were estimated by comparison with the reference substances. For the reference materials of Cd-O, Cd-acetate and pectin-Cd were used, and for Cd-S, phytochelatin-Cd was used.
a) *R* factor: reliability factor for goodness of fit calculated as residual between fit and experimental data.

When the plant was cultured under the condition of 2000 μM Cd, the ratio of Cd compounds accumulated in roots was 72 % as Cd-O and 28 % as Cd-S (Table 2). The proportion of Cd-S was markedly increased in the plants exposed to 2000 μM Cd compared to those exposed to 200 μM Cd. This observation suggests that stronger Cd stress induced an increase in the chelate formation of Cd by the thiol compounds in the underground parts. On the other hand, the ratio of Cd-S in the aerial part was 36 %, and the ratio of the Cd-S abundance was not significantly affected even when it was exposed to the high concentration of Cd (Table 2).

## Discussion

Many ordinary plant species, such as tobacco, often show a significant reduction in their growth even under the conditions containing low Cd concentrations (Misra and Gedamu, 1989). Cd stress in plants occasionally causes ROS induction and results in cell death (Petrov et al. 2015). Glutathione functions as an antioxidant because its thiol group acts as an electron donor on active oxygen to reduce and remove ROS (Pompella et al. 2003). When cells are exposed to increased levels of oxidative stress, glutathione is oxidized to glutathione-S-S-glutathione (GSSG) and accumulates. An increased ratio of GSSG to glutathione is an indication of oxidative stress (Szalai et al. 2009). Phytochelatin, a small peptide composed of multiple units of reduced glutathione, protects cells by chelating harmful heavy metals such as Cd and subsequently localizes and accumulates in vacuoles and cell walls (Cobbett, 2000). Therefore, a loss-of-function transformant of glutathione synthesis becomes strongly susceptible to Cd stress (Howden et al., 1995).

The amount of total glutathione in underground part was decreased, but oxidized glutathione wasn’t detected (Fig. 1a). We have reported that *A. yokoscense* accumulated Cd in the distal roots but less in shoots, and no ROS in the stele of the root were induced when exposed to 0.1 and 100 μM Cd, respectively (Ukai et al. 2020).

These results indicate that *A. yokoscense* suffers little oxidative stress due to Cd exposure. Whereas an increase and accumulation of thiol compounds were observed in the stele of *A. yokoscense* when exposed to 100 μM Cd (Fig. 1b). The thiol compounds that increased in concentration in Cd-exposed *A. yokoscense* are possibly phytochelatins. If the thiol compounds are phytochelatins, the decrease in total glutathione content can be explained by the consumption of GSH as a precursor.

When the *A. yokoscense* sporophyte was exposed to Cd for a long period (two weeks), a large amount of Cd uptake was observed even in the aerial part. *A. yokoscense* exposed to a high concentration (2000 μM) of Cd led to more than 3 mg g^−1^ Cd accumulation in the aerial parts without any significant growth inhibition (Table 1). The amounts of Cd accumulation corresponded to approximately two-thirds of those of Cd detected in roots. These results suggest that *A. yokoscense* has a mechanism for protecting against Cd injury in the case of long-term Cd exposure in addition to mechanisms working to block the entry of Cd.

It is known that Cd absorption is often involved in the role of some metal transporters (Thomine et al. 2000). In rice, Nramp5, an Fe transporter present in the cell membrane, is known to be associated with the uptake of Cd into cells (Ishikawa et al. 2012). *A. yokoscense* exhibited the restricted Cd transport into the aerial parts. We assumed that this may depend on the low Cd absorption by a metal transporter. To know this, we examined the ability of Cd uptake of AyNramp5. In the transformant containing *AyNramp5a*, an *OsNramp5*-like gene whose protein functions as a metal transporter, the amount of Cd absorption was significantly increased (Fig. 2). This result reveals that AyNramp5a could absorb Cd into cells. This also suggests that Cd will be taken into *A. yokoscense* cells via metal transporters such as Nramp5. It is not the nullified transport activities of Cd by metal transporter such as Nramp5 once supposed, but some other mechanisms that achieve the restricted Cd transport to the aerial part. In addition, it is noteworthy that the rice calli overexpressing this gene still maintained high levels of Fe and Mn concentrations under the Cd stress.

The roots of *A. yokoscense* contained significantly more organic acids than those of tobacco (Fig. 3). In mangrove roots, it has been reported that the extracellular secretion of organic acids is involved in resistance to Cd stress (Xie et al. 2013). Sun et al. have shown that the amounts of organic acids and thiol compounds are larger in Cd-tolerant cabbage varieties than in Cd-susceptible varieties. In Cd-tolerant plants, it has also been suggested an increase in thiol compounds and organic acids in the cell wall in the roots where Cd resistance is achieved by localizing Cd (Sun et al., 2013). In this study, the SR-XRF analysis revealed that *A. yokoscense* accumulated a large amount of Cd in roots, in which a large proportion was localized in the cell wall and the intercellular space in the epidermal region (Fig. 4). This suggests that *A. yokoscense* preferentially localized Cd in the cell wall of the roots, suppressing its intrusion. Organic acids in this region may contribute to suppressing the Cd uptake into the cells. In the roots of *A. yokoscense*, the existence of a Casparian strip has been suggested (Ukai et al. 2020). This might restrict Cd intrusion into the central region. Further study using normal fern as the reference is needed to confirm our hypothesis.

The roots exposed to 200 μM Cd contained Cd-O in a large proportion (Table 2). These results suggest that Cd was primarily bound to oxygen in organic acids and consequently localized in the cell wall. This result suggests that root organic acids may be involved in the tolerance mechanism to short-term Cd exposure. In the roots exposed to 2000 μM Cd, the ratio of components of Cd-S molecules increased (Table 2). Cd-S compounds conjugated with glutathione and phytochelatin may function to chelate Cd and detoxify it by localizing it to vacuoles. This observation suggests that strong Cd stress induces the formation of Cd-S in roots to reduce the toxicity in the cell. In the aerial parts, the proportion of Cd-S was approximately 40 % in both plants exposed to 200 and 2000 μM Cd, which was larger than that of Cd-S in roots (Table 2). This suggests that thiol compounds such as phytochelatin are recruited for detoxification of Cd in the cells.

As mentioned above, *A. yokoscense* is suggested to have an excellent system involved in tolerance to Cd intrusion. In the case of Cd exposure at a low concentration and in the short term, Cd-O compounds could be formed by conjugation with organic acids abundantly present in the cell walls of roots, thereby suppressing Cd uptake into the cells. When *A. yokoscense* is exposed to high concentrations of Cd or for long-term exposure, Cd would enter cells via a metal ion transporter and induces the generation of thiol compounds, such as glutathione and phytochelatin, which may contribute to the formation of Cd-S compounds. Cd migration to the aerial part may induce the accumulation of thiol compounds in the stele. These factors are suggested to suppress the migration from the roots to the aerial parts. In the aerial parts, the proportion of Cd-S forms increases due to Cd accumulation; thiol compounds such as phytochelatin may be recruited to work for detoxification of Cd in the aerial cells and lead Cd to localize to vacuoles.

## Data Availability

The dataset of the comprehensive analysis on the *A. yokoscense* complehensive transcripts has been registered by the DNA Data Bank of Japan Sequence Read Archive under accession number DRA008924. Nucleotide sequences of the individual *AyNramp5* genes have been registered by the DDBJ database with the following accession numbers: ICTA01000001 (*AyNramp5a*), ICTA01000002 (*AyNramp5b*), ICTA01000003 (*AyNramp5c*), ICTA01000004 (*AyNramp5d*), ICTA01000005 (*AyNramp5e*), ICTA01000006 (*AyNramp5f*), ICTA01000007 (*AyNramp5g*), and ICTA01000008 (*AyNramp5h*).

## Acknowledgements

We thank Mr. Yuki Yamamoto, Mr. Shunsuke Yanagisawa and Dr. Hiroshi Teramura for technical assistance and Dr. Izumi Nakai and Dr. Shin-nosuke Hashida for valuable discussion.

## Author Contributions

Y. Ukai performed physiological and genetic analysis and wrote the article. These works were carried out in Tokyo University of Science; H. Taoka analyzed the physiological analysis and XRF and XAFS analysis; M. Kamada and Y. Wakui performed physiological and genetic analysis; F. Goto, K. Kitazaki, and T. Abe designed the study and analysis; A. Hokura and T. Yoshihara designed and conducted the study and wrote the article; H. Shimada designed and conducted the study, analyzed the data and wrote the article.

---

**Supplementary Figure S1.**
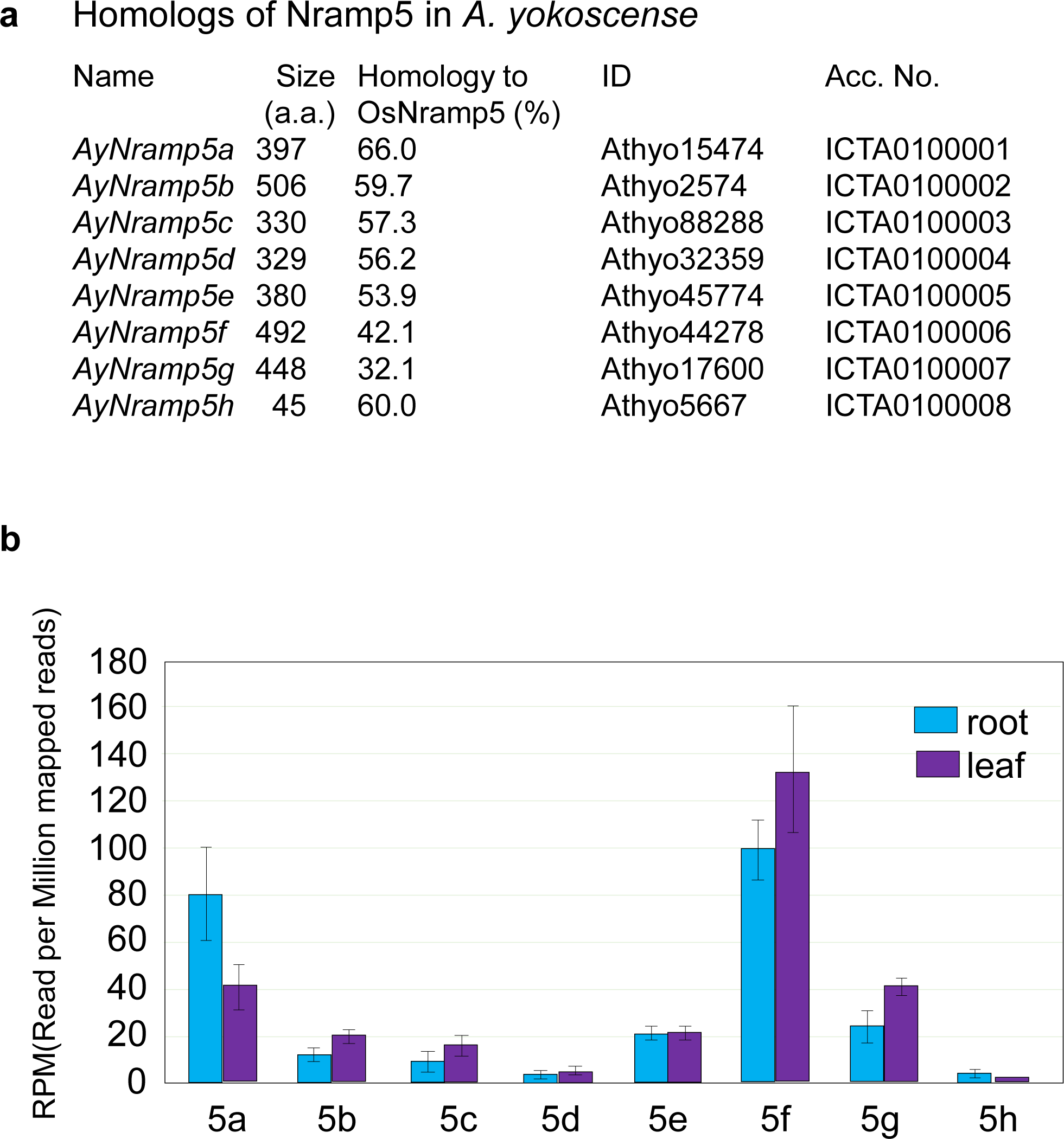
(a) List of *A. yokoscense* Nramp5 genes. “Size” indicates the number of amino acid residues in the registered cDNA. The cDNA of AyNramp5h contains a partial sequence without translational termination. ID and Acc. No. indicates the corresponding ID names in the dataset registered by DRA008924, and accession numbers of the DDBJ database. (b) Suggestive expression levels of the predicted *Athyrium yokoscense Nramp5* genes. Expression levels are represented as the numbers of reads per million mapped reads (RPM), whose original data have been reported previously (Ukai et al. 2020). *Nramp5* genes are indicated by names without the prefix “AyNramp”, such as 5a and 5b.

**Supplementary Figure S2.**
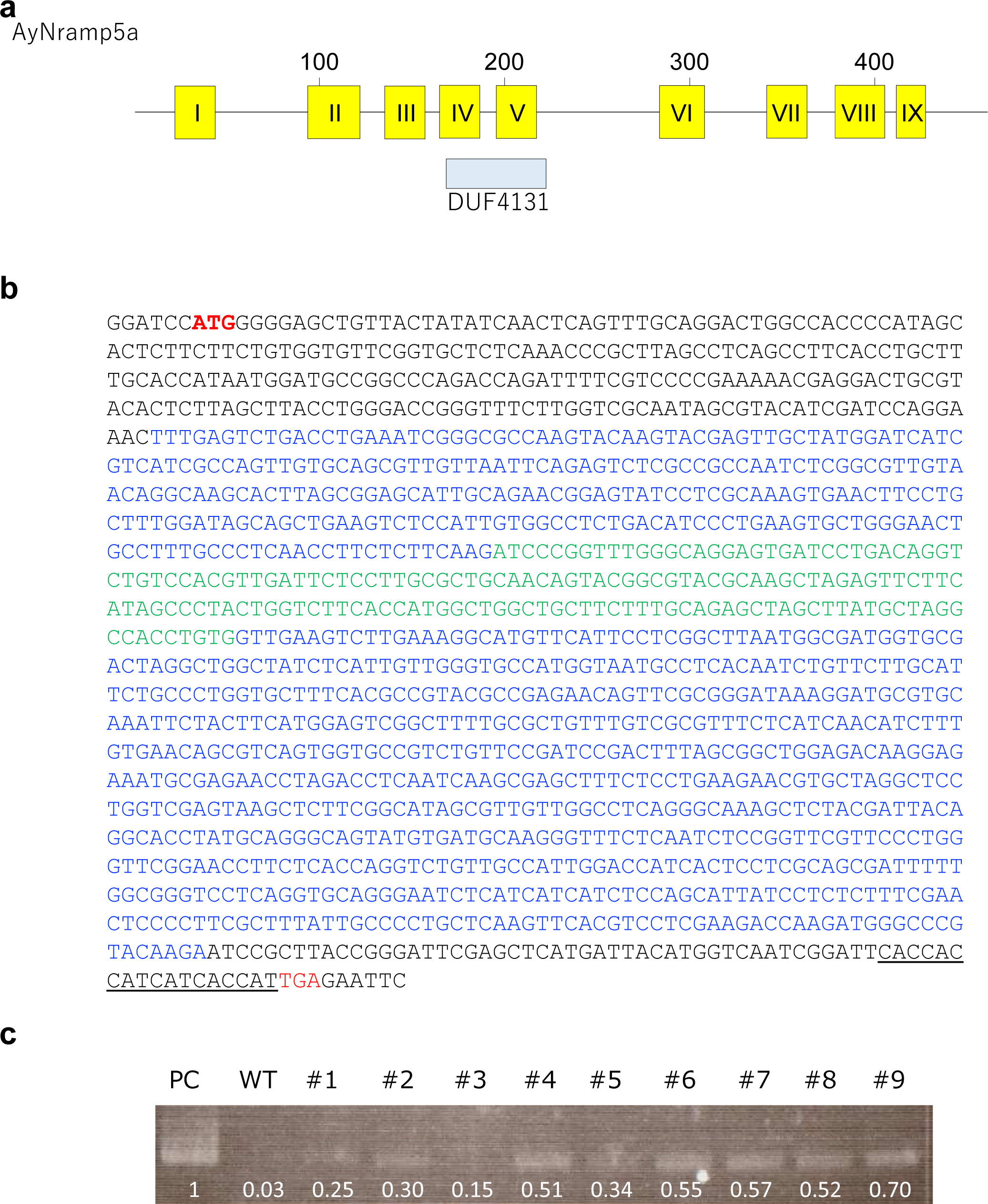
(a) Schematic representation of the structure of AyNramp5a. Yellow boxes indicate the transmembrane domains. The DUF4131 region is indicated by a light blue box. (b) Nucleotide sequence of the chemically synthesized AyNramp5a-6×His gene, which was applied for expression in rice cells. Letters in red indicate the initiation and termination codons, and the sequence of 6×His is underlined. Letters in blue indicate the Nramp domains, and letters in green indicate the region corresponding to DUF4131 domains. (c) Gene expression analysis of AyNramp5a in transgenic plants by RT-PCR. PC:pGWB2—*AyNramp5*—6×His plasmid, WT: wild type, #1 - #9: independent transformant callus line

**Supplementary Table 1.** Analytical conditions of μ-XRF imaging.

| Figure number | X-ray energy /keV | Beam size /$\mu\text{m}$(V) x $\mu\text{m}$(H) | Step size /$\mu\text{m}$(V) x $\mu\text{m}$(H) | Imaging area /$\mu\text{m}$(V) x $\mu\text{m}$(H) | Dwell time /s point<sup>-1</sup> |
| --- | --- | --- | --- | --- | --- |
| Fig. 4a | 30 | 0.85 x 1.6 | 2 x 2 | 122 x 232 | 0.3 |
| Fig. 4b | 30 | 0.85 x 1.6 | 2 x 2 | 252 x 168 | 0.3 |

The detailed conditions shown in Fig. 4 are summarized.

